# Cataloging the potential functional diversity of Cacna1e splice variants using long-read sequencing

**DOI:** 10.1101/2022.04.06.487199

**Authors:** Shamsuddin Bhuiyan, John R. Tyson, Manuel Belmadani, Jordan Sicherman, Terrance P. Snutch, Paul Pavlidis

## Abstract

Voltage gated calcium channels (VGCCs) regulate the influx of calcium ions in many cell types, but our lack of knowledge about the plethora of VGCC splice variants remains a gap in our understanding of calcium channel function. A recent advance in profiling gene splice variation is to use long-read RNA-sequencing technology. We sequenced Cacna1e transcripts from the rat thalamus using Oxford Nanopore sequencing, yielding the full structure of 2,110 Cacna1e splice variants. However, we observed that only 154 Cacna1e splice variants were likely to encode for a functional VGCC based on predicted amino acid sequences. We then computationally prioritized these 154 splice variants using expression and evolutionary conservation and found that four splice variants are candidate functionally distinct splice isoforms. Our work not only provides long-read sequencing of Cacna1e for the first time, but also the first computational evaluation of which Cacna1e splice variants are the best candidates for future follow-up.

**SIGNIFICANCE STATEMENT:** Voltage gated calcium channels (Cacna1x genes) are implicated in many neurological disorders and their encoding genes are predicted to have complex patterns of alternative splicing. Previous approaches relied on short-read RNA-seq to characterize calcium channel splice variants. Here, we use long-read nanopore sequencing to establish a set of Cacna1e transcripts in the rat thalamus and use computational methods to prioritize four transcripts as functionally distinct splice isoforms. Our work to provide the field with prioritized transcripts will not only improve our understanding of Cacna1e function but its role in disease as well.

## INTRODUCTION

Voltage-gated calcium channels (VGCCs) play a crucial role in regulating calcium ion influx in neuronal cells. Mutations in the pore-forming or α_1_-subunit genes that encode VGCCs (CACNA1x, also termed Cavs) are linked to many neurological disorders (Heyes et al., 2015). Despite their importance, our understanding of VGCCs remains incomplete as each gene undergoes extensive alternative splicing. To fully appreciate VGCCs, we must first generate the complete splicing repertoires of each CACNA1 gene and determine those splice variants that impact the overall channel function. Here, we describe our work to profile the distinct splice variants of the rat Cacna1e gene using long-read RNA-sequencing.

Cacna1e encodes for the α_1_-subunit of the Cav2.3 channel (Soong et al., 1993; Williams et al., 1994; Wormuth et al. 2016). Cav2.3 currents uniquely exhibit biophysical and pharmacological characteristics representative of both high-voltage-activated and low-voltage-activated VGCCs. Cacna1e knock-out in rodents caused a decrease in calcium current in rat hippocampal pyramidal cells, an increased sensitivity to inflammatory pain, and a resistance to drug-induced seizures (Saegusa et al., 2002; Simms and Zamponi, 2014; Sochivko et al., 2002, p. 2; Weiergräber et al., 2007; Zaman et al., 2011). Gain-of-function mutations have been associated with epileptic encephalopathy, macrocephaly, and dyskinesia (Helbig et al., 2018; Weiergräber et al., 2006; Wormuth et al., 2016; Zaman et al., 2011). Furthermore, due to the differential expression distribution of splice variants across tissues, CACNA1E potentially has tissue-specific functions (Fang et al., 2007).

The extent to which alternative splicing expands genomic functional diversity remains the subject of debate, and CACNA1E, with its extensive list of splice variants, is a notable example (Bhuiyan et al., 2018; Blencowe, 2017; Tress et al., 2017). Like other VGCCs, the large Cacna1e gene contains at least 36 exons. The a1-subunit must be at least 2000 amino acids long and contain four pore-forming domains to make a functional pore (Catterall, 2011) and mutations in any of these pore-forming domains can result in a dysfunctional channel (Guida et al., 2001). The multiple splice variants for CACNA1E generally have similar exons encoding for the conserved pore-forming domains; however they vary in their use of exons in between the pore-forming domains (the linker regions) and carboxyl terminal regions (Lipscombe et al., 2013b; Lipscombe and Andrade, 2015).

When examining eight CACNA1E “alternative” splice variants, Pereverzev and colleagues found that different variants encode for channels with distinct inactivation and recovery time courses (Pereverzev et al., 2002). Additionally, Klockner and colleagues noted distinct binding affinities among Cav2.3 regulatory proteins and CACNA1E splice variants (Klöckner et al., 2004). Given CACNA1E antagonists (e.g. Topiramate) are used to treat to seizures, understanding the role of CACNA1E splice variation towards developing better drug targets as different splice variants have different pharmacological sensitivities (Kuzmiski et al., 2005; Lipscombe and Andrade, 2015).

Short-read sequencing (~150-175 bp) has been previously employed to characterize CACNA1E splicing profiles, which is limited by the inability to resolve the relationship between an individual short read and the full length transcript structure (Lipscombe et al., 2013a; Steijger et al., 2013). An alternative to short-read sequencing, long-read nanopore-derived RNA-seq data using Oxford Nanopore Technology (ONT) technology provides the opportunity to define the entire splice variant repertoire (Clark et al., 2019). Clark and colleagues used MinION sequencing to establish CACNA1C transcriptional complexity in the human brain (Clark et al., 2019). They reported only a trivial minority of the CACNA1C splice variants were previously included in GENCODE and speculated that these newly identified variants contribute to the functional diversity of CACNA1C in different brain regions.

The lack of a comprehensive CACNA1E splice variant catalog likely impacts the various genomic databases that report splice variants such as Ensembl, and highlights the previously noted disconnect between computational tools and the experimental literature (Bhuiyan et al., 2018). Bhuiyan and colleagues previously reported the failure to identify splice variants in Ensembl for about a third of genes with literature evidence for functionally distinct splice isoforms (FDSIs) (Bhuiyan et al., 2018). This disconnect likely impacts the ability to computationally evaluate alternative splicing in genes with complex splicing patterns.

In this study, we characterized the splicing profile for rat Cacna1e using ONT RNA-seq data. We identified the structures and splice junctions for 2,110 Cacna1e splice variants. Using computational annotations, we established a putative set of Cacna1e splice variants. In doing so, we have demonstrated the potential functional diversity of these genes while maintaining a more accurate characterization of the splice variant structure than previously available. This improved transcript catalog can aid computational tools and provide experimentalists potentially interesting splice variants for investigation.

## METHODS

### Targeted amplification for five α_1_-subunit genes of GAERS and NEC rats

We performed targeted amplification of Cacna1e transcripts in GAERS and NEC (Non-Epileptic Control) rats. Total RNA was extracted from the thalamus of 2 GAERS (10 and 90 days) and 2 NEC (10 and 90 days) male rats using a MagMax Kit (Ambion) and full length cDNAs generated using SuperScriptII with oligo-dT priming (Invitrogen). These samples will be referred to as GAERS10, GAERS90, NEC10 and NEC90. Gene specific amplicons for Cacna1e genes (Forward: ATGGCTCGCTTCGGGGAG; Reverse: CTAGCACTTATCGTCTTCTTC) was generated using PCR with the Elongase enzyme (Invitrogen). PCR products were then gel purified using a gel extraction kit (QIAGEN) and DNA eluted in TE and stored at −20°C ready for sequencing. All animal procedures were performed in accordance with the [Author University redacted] animal care committee regulations.

### ONT MinION sequencing of amplicons

The generated gene specific amplicons were processed as per the Oxford Nanopore Technologies (ONT) SQK-LSK108 adapter ligation procedure. In brief, the DNA amplicon molecules were treated to generate A-tailed molecules allowing the ligation of the ONT specific adapter onto the amplicon ends. These adapted molecules were then run on the ONT MinION device and signal data captured for base calling over a 48hr period. Sequence data was generated from the raw captured data using ONT specific software (Guppy 3.4) on a GPU enabled desktop PC. Raw read data are available [link redacted].

### Short-read sequencing of genes and alignment

The generated gene-specific amplicons were short-read sequenced at the BC Genome Sciences Center and data was provided back in SOLEXA format. Raw short-reads will be available through the Short Reads Archive (SRA) [accession redacted]

We reprocessed the rat transcriptomic SOLEXA data. Since these reads were in SOLEXA format, we converted them into FASTQ format in order to run them through our short-read RNA-seq pipeline.

The rat transcriptome reference was prepared using the “rsem-prepare-reference” script provided by the software package “RNA-seq by expectation-maximization” in RSEM (Li and Dewey, 2011). The assembly version used was Ensembl Rnor6.0, obtained through Illumina for the iGenomes collection. (https://support.illumina.com/sequencing/sequencing_software/igenome.html). Short-reads were processed as single-end (no mate pairs) and aligned using the STAR aligner (Dobin et al., 2013) version 2.4.0h. provided as input to the quantification scripts from RSEM v1.2.31. Default parameters were used (with the exception of parallel processing and logging related options). The count quantification matrix of splice junctions (SJ.tab.out) was used for downstream FLAIR analysis.

### Determining splice variants using long read RNA-seq data and FLAIR

We aligned our long-reads for each sample to the Ensembl rat genome (rn6) using FLAIR (Tang et al., 2018) and minimap2 (Li, 2018). With the addition of the splice junctions found in the Illumina data, FLAIR corrected the reads in each alignment file. In short, this means that any novel splice junctions are merged to an existing splice junction if the junction is within a 10-nucleotide window of the existing junction. Finally, FLAIR collapsed any reads having the same transcription start site and same splice junctions across all samples into a single splice variant. FLAIR removed any splice variants without support from at least 3 reads.

### Definition of functional Cacna1e splice variant

A gene with functionally distinct splice isoforms (FDSIs) is defined as a gene with two or more splice isoforms that are necessary for the gene’s overall function. Under this definition, each individual isoform must cause a change in phenotype after experimental depletion (Bhuiyan et al., 2018). For the purposes of computationally characterizing a candidate FDSI from Cacna1e splice variants, we define a candidate Cacna1e FDSI as “a splice variant evolving under selection (conservation) with all 4 pore-forming domains necessary for calcium passage into the cell”. The requirement that the splice variant has all 4 pore-forming domains implicitly means that the splice variant must have an open reading frame (ORF) of at least 6000 bp (2000 amino acids). Conserved splice variants indicate that the splice variant is necessary for reproductive success and sequence conservation acts as a proxy for functional importance.

### Cacna1e splice variant annotations

Our annotation pipeline is available on [git link redacted]

We subsetted the 6,252 splice variants from FLAIR to 2,110 Cacna1e splice variants and then reduced all splice variants into their exons and identified exons shared between splice variants. Further, we annotated any exon that was not entirely found in Ensembl as novel. We then annotated all splice variants with the number of MinION reads which support it (Supplementary File 2).

As many transcripts are lowly expressed, we consider genes with multiple appreciably expressed splice variants as stronger candidates for having FDSIs (Gonzalez-Porta et al., 2013). We annotated each Cacna1e splice variant with a gene-specific ranking and gene-specific expression ratio. The most highly expressed splice variant was ranked ‘1’, the second most expressed splice variant was ranked ‘2’, and so on until all splice variants were ranked. We produced ranks based on the total expression across all samples. Using the gene-specific rankings, we calculated an expression ratio between each Cacna1e splice variant and Cacna1e’s rank 1 (most expressed) splice variant. For the purposes of prioritization, we considered splice variants where the expression level of the splice variant was at least 50% of the expression level of the rank 1 splice variant (expression ratio less than 2).

We annotated all splice variants and their individual exons with their average PhastCons (Siepel et al., 2005) and PhyloP (Pollard et al., 2010) basewise scores from a 20 vertebrate species alignment, downloaded from the UCSC genome browser for build rn6 (Rosenbloom et al., 2015). The average score for each exon was calculated using Kentutils. In order to determine if there was any selection pressure upon the splice sites, we also annotated the average PhyloP conservation of the two intronic bases next to the 5’ side of the exon and the two intronic bases next to the 3’ side of the exon.

Using TransDecoder (https://github.com/TransDecoder/TransDecoder) we predicted translation products of the 2,110 splice variants from Cacna1e for any ORFs larger than 2,000 amino acids with a start codon. These ORFs do not necessarily contain a stop codon. We then queried the Conserved Domain Database (CDD) with the translated sequences and annotated each splice variant with the protein domain hits the database returned (Marchler-Bauer et al., 2015).

We then performed homology searches against the GenBank nr database (Pruitt et al., 2007) on all exons using NCBI’s tBLASTn (Camacho et al., 2009). First, we extracted the sequences for all alternatively spliced exons. Each alternatively spliced exon’s sequence was concatenated with the sequences of its flanking exons. The set of three exonic sequences was BLAST-ed against the human, mouse, rat, zebrafish, fugu, coelacanth, lamprey and spotted gar data in the nr database. BLAST results were next filtered using an e-value threshold of 0.0001, gap threshold of less than 30%, query coverage threshold of 80% or more, and a percent identity threshold of 30% or more.

### Visualization tool

We developed a visualization tool using R that uses three input files: a BED-formatted file of all splice variants (FLAIR output), a tab-delimited CDD output file, and a CSV formatted file of each splice variant’s expression. The visualization tool calculates, based on chromosomal position and exon sizes, the set of similarly annotated exons that are preserved between all splice variants of interest, and draws the result as a stack of aligned exon patterns. Since introns were disproportionately large as compared to the exons present in Cacna1e splice variants, drawing lengths of introns were made arbitrary. The tool and example inputs are available online at [git link redacted].

## RESULTS

We profiled the Cacna1e transcriptome using RNA isolated from the rat thalamus for targeted MinION cDNA sequencing. Our analysis encompassed 4,060,847 reads and initially yielded 2,110 potential splice variants (Table 1 and 2; Supplementary File 1 and Supplementary File 2).

**Table 1:** Summary of targeted MinION RNA-seq data. All four samples were sequenced with gene specific PCR primers. Column headings: NEC 10 – Non-epileptic control at 10 days, NEC 90 – Non-epileptic control at 90 days, GAERS 10 – Genetic Absence Epilepsy Rat from Strasbourg at 10 days, GAERS 10 – Genetic Absence Epilepsy Rat from Strasbourg at 90 days.

|  | NEC 10 | NEC 90 | GAERS 10 | GAERS 90 |
| --- | --- | --- | --- | --- |
| # Reads | 3,514,669 | 4,958,824 | 3,255,180 | 6,127,231 |
| # Bases | 20,259,569,128 | 23,582,737,080 | 19,092,042,421 | 31,856,992,171 |
| Unmapped reads | 31,356 | 31,356 | 17,311 | 5,773 |
| Mean length of reads | 5,764 | 4,756 | 5,865 | 5,199 |
| Median length of reads | 7,952 | 8,948 | 8,134 | 7,902 |
| Number of reads FLAIR assigned to splice variants | 3,494,983 | 4,881,246 | 3,243,064 | 6,076,453 |
| Number of reads removed | 19,686 | 77,578 | 12,116 | 50,778 |
| Mean read quality | 10.50 | 9.80 | 9.90 | 9.90 |
| Median read quality | 10.80 | 9.90 | 10.20 | 9.70 |

**Table 2:**
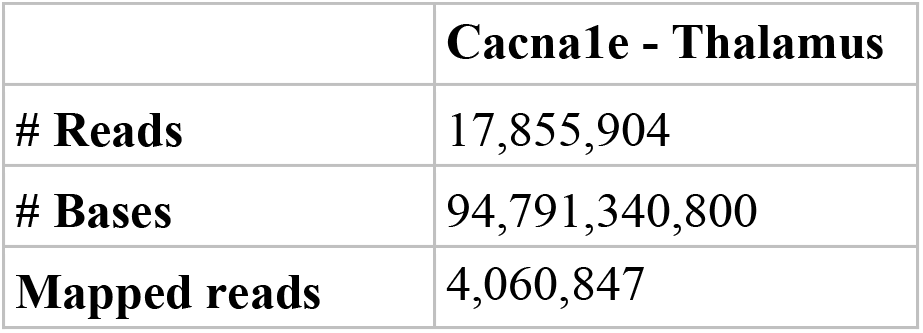
Summary of short-read RNA-seq data.

|  | Cacna1e - Thalamus |
| --- | --- |
| # Reads | 17,855,904 |
| # Bases | 94,791,340,800 |
| Mapped reads | 4,060,847 |

The goal of this analysis was to identify which splice variants could likely form functional calcium channels and which are most likely to contribute to Cacna1e’s functional diversity.

### Detection of candidate Cacna1e splice variants

Data were initially filtered for reads that mapped to Cacna1e, yielding an initial pool of 2,110 variants for Cacna1e that passed quality control (see Methods). Approximately 53% of the reads were at least 6,000 nucleotides with a median length of 5,725 nucleotides. The number of reads that mapped to the same splice variant was used as a proxy for the splice variant expression level. We observed a moderate correlation (Spearman r = 0.59) between splice variant size and the number of supporting reads per splice variant.

We determined how many of the a1-subunit splice variants had the translational potential for a functional a1-subunit (Table 3, Figure 1). Only 238 of the 2,110 Cacna1e splice variants that could translate to a protein longer than 2,000 amino acids (all Transdecoder predicted amino acid sequences provided in Supplementary File 3). Furthermore, even fewer of these splice variants are likely to have an ORF with all four pore-forming domains. Based on protein domain annotations from CDD, only 154 of the 238 Cacna1e splice variants had an ORF longer than 2000 amino acids that span sequences for the four pore-forming domains (full CDD results in Supplementary file 4 and Supplementary File 5).

**Table 3:**
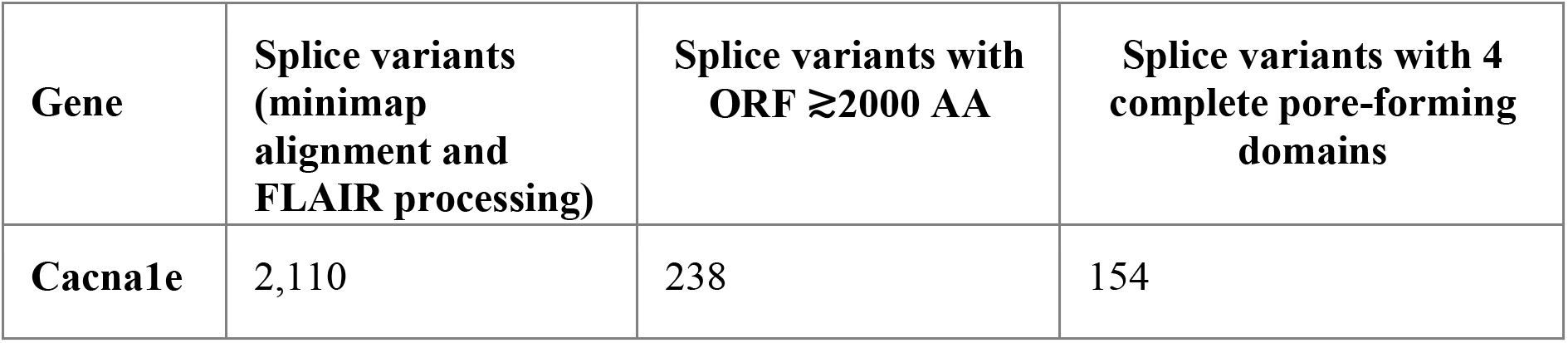
Summary of putative splice variants detected in long-read RNA-seq data. Initial set of Cacna1e splice variants was 2,110 splice variants. Of those 2,110 splice variants, 238 contained an open reading frame of 2,000 amino acids or greater. We further filtered these splice variants to 154 based on whether they contain the four necessary pore-forming domains.

**Figure 1:**
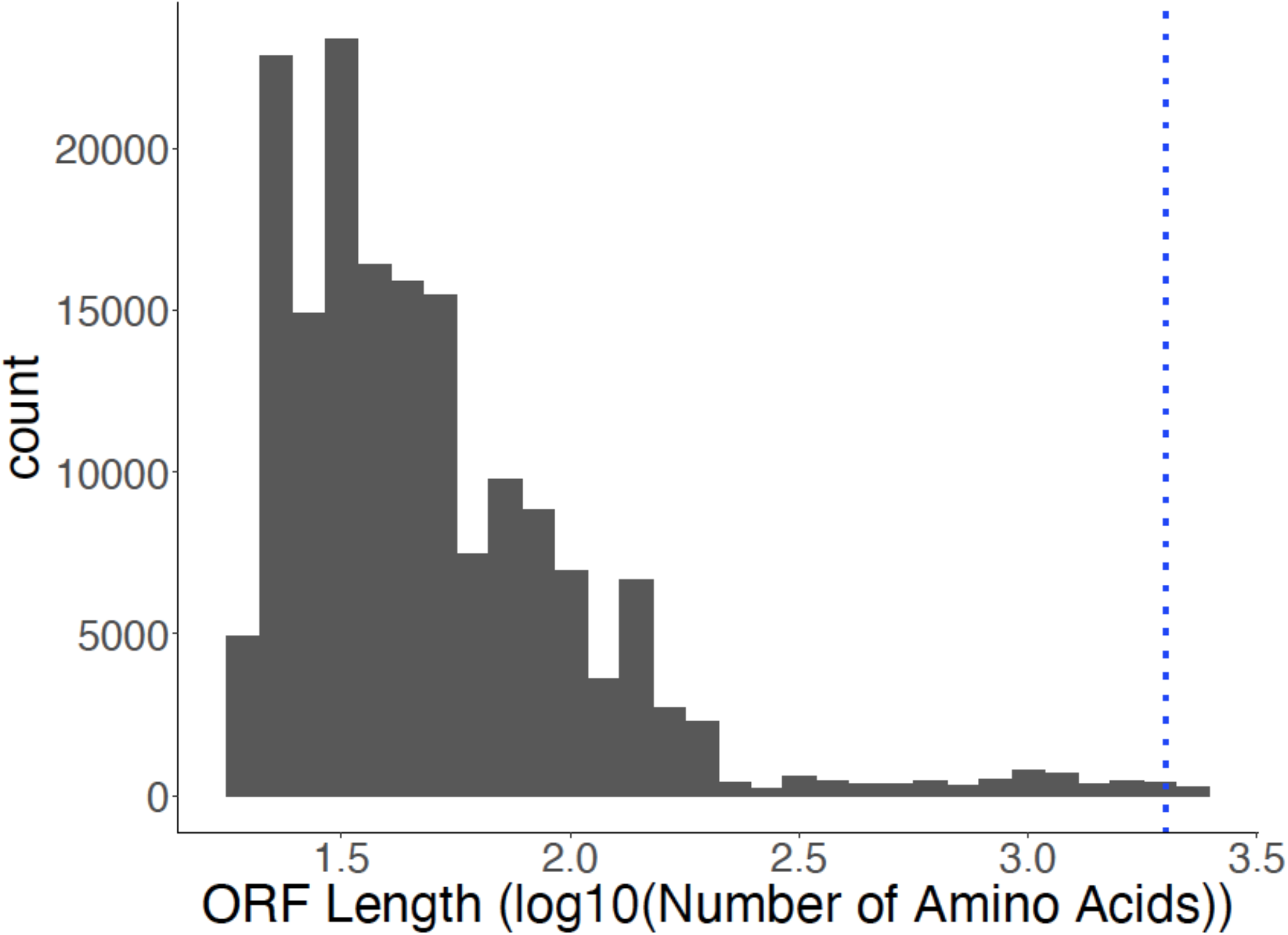
The majority of detected splice variants did not contain an open reading frame (ORF) of at least 2,000 amino acids. ORFs were defined as containing a start codon, but not necessarily a stop codon. The X-axis shows the distribution of log10(ORF) lengths. The dashed blue line indicates the minimum size expected for a functional channel.

### Expression profiles of Cacna1e splice variants

In our set of putative Cacna1e splice variants, we detected the splice variant characterized by Soong and colleagues (Soong et al., 1993). None of the transcripts in our long-read data matched the transcripts reported in Ensembl v98, including the sole Ensembl transcript that codes with an ORF over 2000 amino acids (ENSRNOT00000003928). This remained the case even after inspecting the MinION data at the raw read level to ensure that data relevant to this transcript had not been filtered out by our processing pipeline.

Using the number of reads mapped to each splice variant as a proxy for splice variant expression, we observed that the top four splice variants expressed at similar levels (Figure 2). These four splice variants each had an expression ratio of less than 2, which indicated that the rank 1 (most expressed) splice variant was double the expression of all but three splice variants (Figure 2A). The rank 1 Cacna1e splice variant matches the splice variant structure and protein sequence of Soong et al. (Soong et al., 1993). We mapped 498,616 reads to this most abundantly expressed splice variant, while the second most abundantly expressed splice variant had 435,458 reads. The top three splice variants contributed similarly to the total Cacna1e expression among the 4 samples: 12.2%, 10.7%, 8.8% and 6.5% (Figure 2B and 2C). However, when stratifying number of reads per splice variant by sample, we observed that the Soong et al. splice variant is the most abundant splice variant in our GAERS90 and NEC90 samples, and fifth most abundant in GAERS10 and NEC10 (Figure 2D).

**Figure 2:**
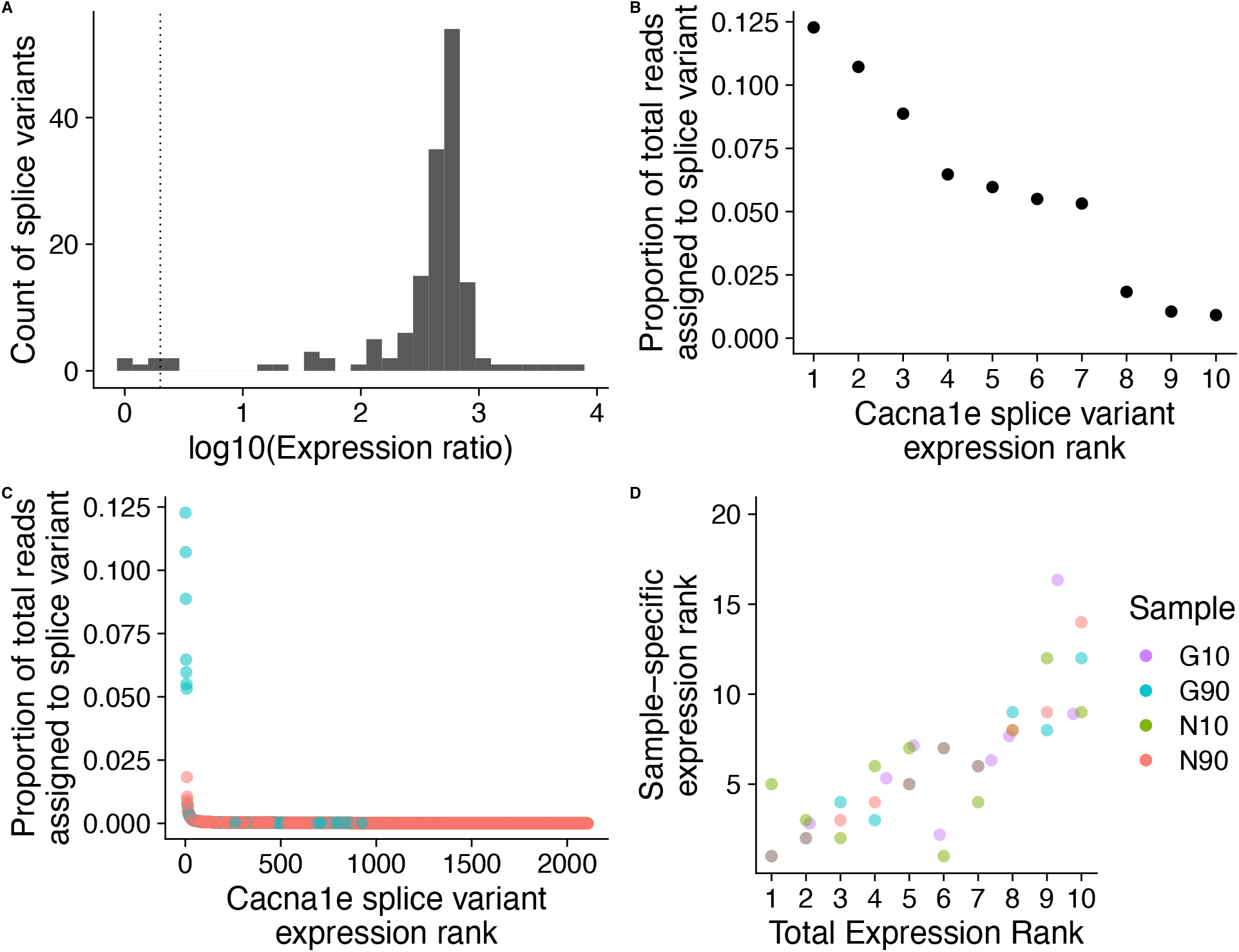
Highly expressed splice variants account for 12.5% of Cacna1e gene expression. A) *Four splice variants have an expression ratio of less than 2.* Each splice variant’s expression ratio is calculated by the most expressed splice variant’s expression divided by the splice variant’s expression. The dotted line indicates where the expression ratio would equal 2. B) *Top 10 most expressed Cacna1e splice variants pooled across all samples.* The X-axis shows the rank of the splice variant based on expression. The Y-axis shows the proportion of the splice variant relative to Cacna1e’s total expression. C) *Contribution of all 2,110 Cacna1e splice variants to the gene’s overall expression.* The X-axis shows the rank of the splice variant based on expression. The Y-axis shows the proportion of the splice variant relative to Cacna1e’s total expression. Of the 2,110 variants, the 154 (cyan) have 4 annotated pore-forming domains and account for 61% of Cacna1e’s expression. We did not predict the remaining splice variants to encode for all four pore-forming domains (red) D) *Cacna1e splice variant expression level ranks in each sample*. X-axis show expression rank for splice variant summed across all 4 sample, while Y-axis shows the splice variant expression rank for each sample. Sample is indicated by color. G10 is Genetic Absence Epilepsy Rat from Strasbourg at 10 days, G90 is Genetic Absence Epilepsy Rat from Strasbourg at 90 days, N10 is non-epileptic control at 10 days and N90 is nonepileptic control at 90 days. Example: the splice variant at expression rank 1 summed across all 4 sample is the most expressed splice variant G10 and N10. However, for samples G90 and N90, that same splice variant is at rank 4 and 5 respectively.

In order to assess the potential functional diversity of Cacna1e splice variants, we downloaded the protein sequence for Ensembl Cacna1e (ENSRNOP00000003928) and the Cacan1e splice variant characterized by Soong et al. (1993) then annotated protein domains as a guide for analyzing our Cacna1e splice variants (Figure 3). Both sequences contained the four pore-forming domains necessary for voltage sensitivity and calcium influx. Furthermore, the Ensembl sequence and the Soong et al. sequence contained the GPHH and IQ domains (GPHH – pfam16905 and Ca_chan_IQ – pfam08763 respectively) necessary for Ca-calmodulin regulation of channel activity (Van Petegem et al., 2005).

**Figure 3:**
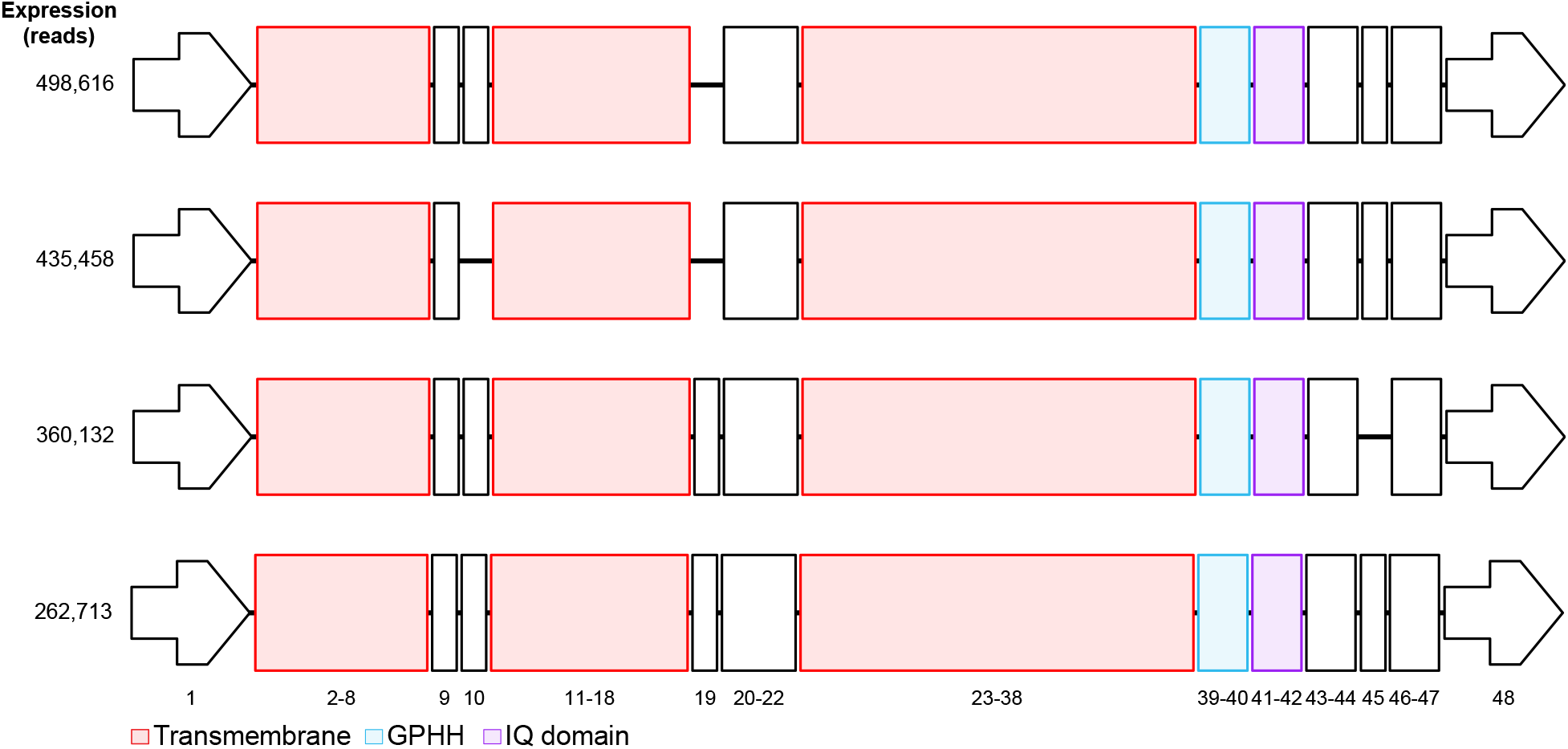
Structure of top 4 most abundantly expressed splice variants. Red indicates a pore-forming domain annotation from CDD (Ion_trans – pfam00520). Blue and purple are calmodulin-binding domain annotation from CDD (GPHH – pfam16905 and Ca_chan_IQ – pfam08763, respectively). Along the side are the number of reads for each splice variant, and long the bottom are the exon numbers. A similar visualization for all 154 splice variants that contained 4 pore-forming domains is provided in Supplementary File 7.

### Cacna1e splice variants contain a conserved cassette exon 19

We further prioritized Cacn1e’s 154 splice variants based on our splice variant-level conservation and expression annotations. We detected 31 cassette exons in our 154 Cacan1e splice variants, and we show where the six most common skipped exons are located relative to channel structure (Figure 4). The three most commonly occurring cassette exons in Cacna1e splice variants were: skipping of exon 10 (40% of reads), insertion of exon 19 (40% of reads), and skipping of exon 44 (45% of reads).

**Figure 4:**
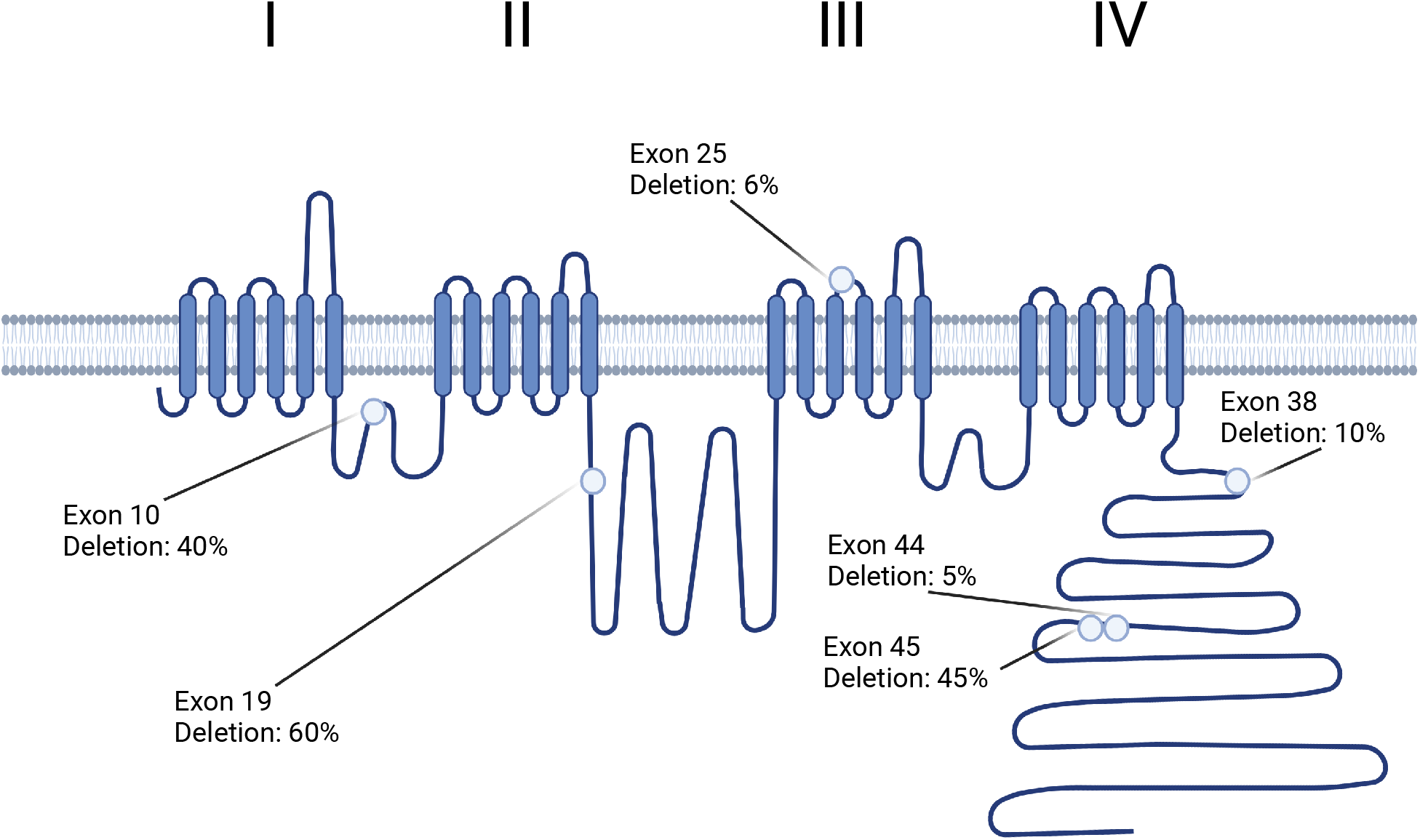
Location of most common splicing events on Cacna1e structure. Here we show where each of these six cassette exons would likely impact the channel, and the percentage of reads that do not contain the exon. Figure created with Biorender.com.

The protein sequences of exons 10, 19 and 44 are all highly conserved with a PhyloP score over 1.3 (Figure 5; PhastCons scores displayed in Extended Figure 1-1). In order to determine whether exon skipping is an inherent aspect of function of the gene or due to splicing errors. We investigated whether the exclusion of these exons was conserved; that is, whether transcripts lacking these exons are present in other species. We performed a BLAST analysis checking whether the protein sequences for each exon together with their flanking exons (i.e. exons 9-10-11, exons 18-19-20, and exons 45-46-47) were present in transcribed sequences in other species. We found evidence of conservation for exon 19 and exon 44 within the jawed-vertebrate phylogeny (fish, rodents and human), but only evidence of conservation for exon 10 skipping within the rodent phylogeny. These are consistent with the potential for exon skipping having a conserved function, although as we discuss later the interpretation is not straightforward due to the potential for conservation artefacts.

**Figure 5:**
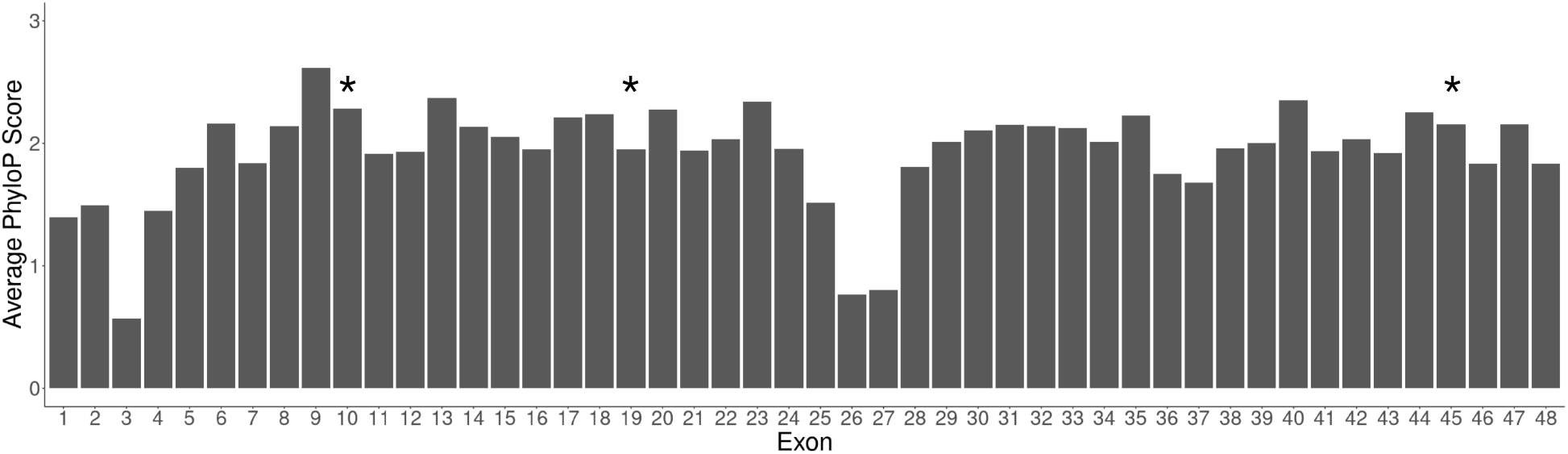
Most Cacna1e exons are well-conserved throughout the vertebrate phylogeny. The UCSC genome browser provides base-specific PhyloP scores. For each exon of Cacna1e, we annotated the average PhyloP score. Our analysis focused on prioritized splice variants with cassette exons 10, 19 and 45 (asterisk).

### Novel Cacna1e splicing events

The 154 Cacna1e splice variants that had the translation potential to encode functional VGCCs contained a total of 31 splicing events (in various combinations), only six of which had previous reports in the literature. In total, these splicing events affect less than 11% of the 154 splice variants (17 splice variants), and the splice variants that contained these splicing events accounted for less than 10% of the total reads. We investigated whether there was conservation of these novel splicing events and whether there was any impact to the protein domains. Sixteen of the splicing events involve exons in a pore-forming domain; however, they did not affect the CDD annotation for the pore-forming domain. Two splicing events are the cassette exons encoding for the GPHH and IQ domain. The seven remaining splicing events are the exons in the N- or C-termini, or in the linker regions.

One subclass of variants (14/153) is predicted to result in the lack of an intact calmodulin-binding domain. The most expressed of these had a total of 292 reads and ranked the 350^th^ most abundant splice variant of the 2,110 Cacna1e splice variants. The most expressed splice variant lacking a complete calmodulin-binding domain contributed to ~0.02% of Cacna1e’s total expression (832 reads). Notably, many of these splicing events lacking calmodulin binding sites have been identified in other species: one is conserved across jawed vertebrates, nine conserved within mammals, and two conserved only within a rodent phylogeny (full results in Supplementary File 6). Given the low estimated expression levels of these 14 splice variants, additional work would be needed to establish their functional relevance.

## DISCUSSION

In this paper, we describe the splicing repertoire for the voltage-gated calcium channel gene (VGCC) Cacan1e using novel targeted long-read RNA-seq data from the thalamus. We then computationally prioritized transcripts in terms of their potential to impact Cacna1e’s functional diversity. This study represents a necessary step towards bridging the gap between claims that alternative splicing vastly increases the functional diversity of VGCC genes and the evidencebased reality of noisy splicing. Though we focused on a specific gene, the approaches we used to computationally assess the biological relevance of splice variants would be applicable to many genes, helping to bridge the gap between raw measures of transcriptional diversity to estimates of functional diversity.

While we detected 2,110 different splice variants for Cacna1e, our analyses suggest that the large majority of these are unlikely to be biologically relevant. In particular, only 157 (7%) are predicted to encode a functional α1-subunit. Of the remaining 93%, only 1,273/1,965 (~68% of the total) have an open reading frame of at least 30 amino acids. The most expressed of these 1,965 splice variants accounts for ~1.8% of the gene’s total measured expression (Figure 2C). While many Nanopore studies have reported expression values by raw reads, we were curious whether library size normalization would influence prioritization, so we quantified the data in terms of transcripts per million reads (TPM; values provided in Supplementary File 2). The raw counts positively correlated with the TPM values of each Cacna1e splice variant (Spearman r = 0.999), and the top ten most expressed splice variants remained at the same rank. The use of TPM values included only two more splice variants in addition to the four splice variants prioritized with an expression ratio of less than 2.0.

The observation of splice variants that do not encode functional products agrees with extensive evidence that RNA splicing is imprecise, such that much “biochemical noise” is produced and which can be captured by modern sensitive molecular biology methods even when present at low levels (Pickrell et al., 2010; Saudemont et al., 2017; Tress et al., 2017; Zhang et al., 2009). While we cannot formally exclude the possibility that some of the non-channel-encoding variants have a function, for follow-up studies we feel it is appropriate to prioritize splice variants for which the most plausible case for functionality can be made.

A further distinction of interest is among splice variants that have *different* and required functions from one another, what we call FDSIs. Computational analyses cannot establish FDSIs, but we applied methods that provide a prioritization. While we detected 154 “full-length” splice variants for Cacna1e, evidence from domain analysis, expression and homology focuses our attention on a subset of four splice variants involving cassette splicing events for exons 10, 19 and 45 in various combinations. We believe the functional consequences of these three exons are worthy of follow-up. Of note, we hypothesize that computational tools are likely to “miss” predicting prioritized splice variants. While Ensembl reports these three exons, none of the Ensembl Cacna1e transcripts match our 2,110 splice variants.

We failed to find any literature discussing the functional effects of exon 10 and our BLAST results mapped to a direct GenBank submission without any associated studies (Human NCBI Reference Sequence: XM_017002244.1). The splicing of exon 10 impacts the linker region between domains I-II, the region of the channel where the β-subunits (Cacnb1 genes) interact with the α1-subunit (Cacna1 genes) to both chaperone the VGCC to the cell membrane and affect channel biophysical parameters (Gonzalez-Gutierrez et al., 2010). Our interpretation of whether the skipping of exon 10 impacts channel function remains limited without *ex silico* validation, but one interpretation may be that the loss of Cacna1e’s exon 10 changes how the calcium channel is chaperoned to the cell membrane.

Both Williams et al. and Schneider et al. reported the cassette splicing of exon 19 (Schneider et al., 1994; Williams et al., 1994). Williams and colleagues showed that splice variants that contained exon 19 changed channel inactivation kinetics, however details of the Schneider et al. study were unavailable. The effects of splice variants that contain exon 19 on whole cell current were later characterized in HEK293 cells and a human pancreatic cell line (Pereverzev et al., 2002; Vajna et al., 2001). Of note, investigating the alternative splicing of a different, but functionally similar VGCC gene, Cacna1b, Gray and colleagues reported that Cacna1b and Cacna1e both had an alternatively spliced 19^th^ exon (Gray et al., 2007). Given the conservation of exon 19 and the location of this exon between domains II and III in both Cacna1b and Cacna1e, Gray and colleagues hypothesized that the exon has an important regulatory role concerning channel function. RT-PCR showed that exon 19 containing transcripts were more prevalent during fetal development than postpartum, and that expression was distinctly absent in the peripheral nervous system (Gray et al., 2007). Our results provide additional transcript structural context for exon 19 which will be useful for future follow-up (Supplementary Figure 6).

We failed to find literature discussing the splicing out of exon 45, and our BLAST results mapped to direct GenBank submissions without any associated study (Human GenBank: AH009158.2, AL161734.12, NG_050616.1). However, exon 45 has been reported to contain *de novo* variants causing epileptic encephalopathy (Helbig et al., 2018). Given the potential role of exon 45 in disease and the strong conservation of the exon 45’s cassette splicing, exon 45 may be interesting for follow-up.

Our study has some important caveats and limitations. Though others have used PCR-based strategies to estimate expression level, the nanopore-based strategy employed here to isolate Cacna1e splice variants is not necessarily expected to lead to accurate expression level estimates for different splice variants, thus relative abundance estimates should be viewed with some caution (Clark et al., 2019). In addition, the use of PCR primers targeting the ends of the only annotated Cacna1e ORF means that we would not detect splice variants containing potential alternative translation start and stops. Another limitation is that we only studied the thalamus, and the splice variants in other brain regions or tissues could be different. Furthermore, with only one sample per condition, and the potential for bias in quantification in cDNA samples which were PCR amplified, we were unable to confidently assess differences in splice variant expression in the GAERS model. Finally, while we used conservation of exon skipping events as a way to evaluate the potential for functional relevance, there is an important caveat in that splicing is known to be imprecise in all species (Melamud and Moult, 2009; Wan and Larson, 2018). As such, we cannot exclude the possibility that observation of an exon skipping event in another species transcriptome is simply the result of chance capture of an “erroneous” isoform. This is made clearer by the observation of “conserved” splicing events that would result in a nonfunctional channel. Given these limitations, we view our study as prioritizing a set of putative FDSIs for Cacna1e, with further high-throughput and low-throughput investigations required to establish their roles and functional importance.

## CONCLUSION

Our work with Cacna1e demonstrates the importance of providing a gene-specific splicing profile using targeted long-read RNA sequencing. We provide transcript structures for 2,110 Canca1e splice variants previously missing in genomic databases. Furthermore, our prioritization using splice variant expression and conservation helps point the field towards potentially interesting splice variants. For Cacna1e, the prioritization using expression and conservation provided a mixture of splice variants that have been functionally investigated in the literature (e.g splice variants containing exon 19) and splice variants without any literature validation (e.g splice variants containing exon 10). The splice variants detected will provide a foundation for future research into VGCCs for both basic research and therapeutic studies.

## Extended data

**Extended Figure 1-1:**
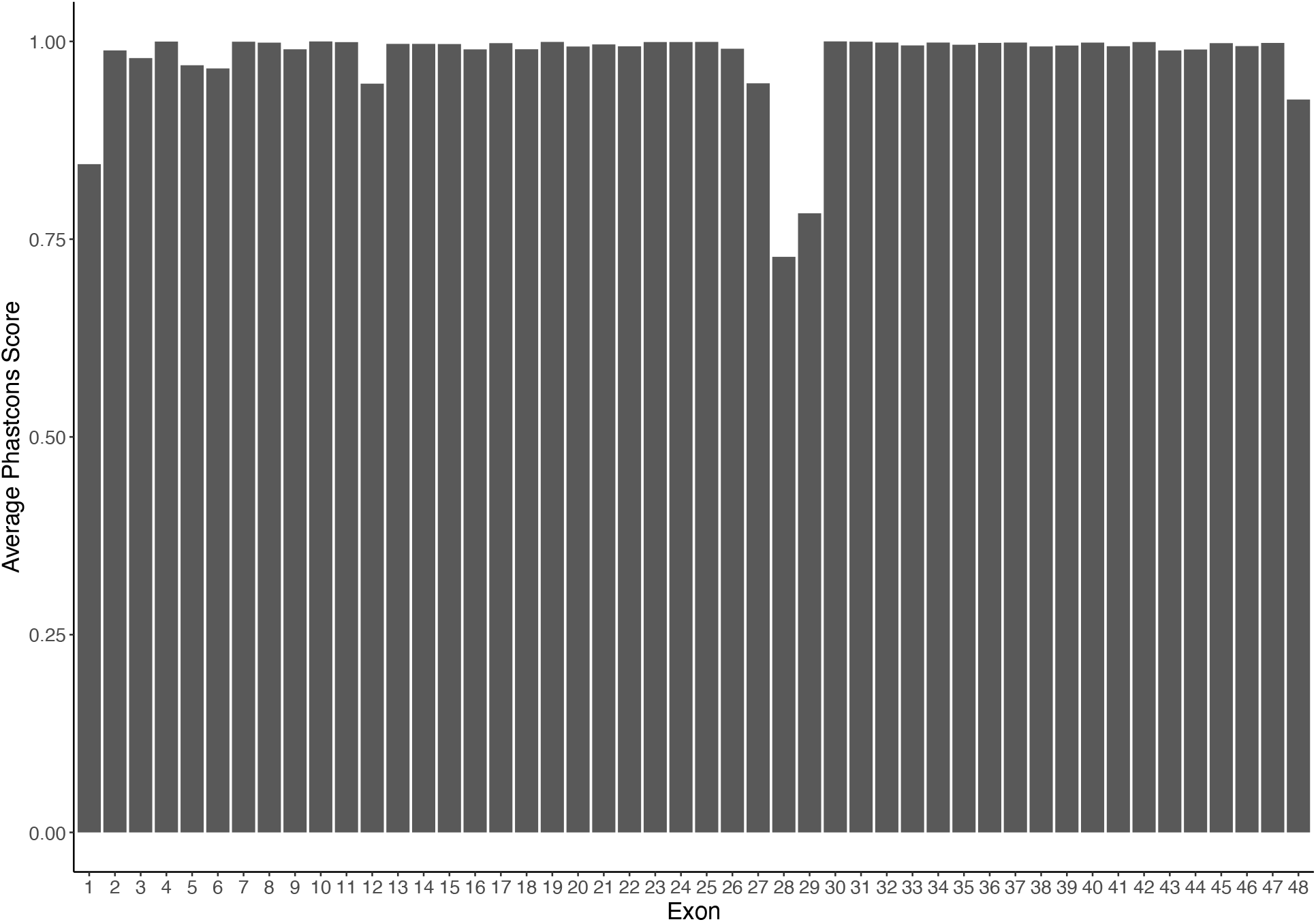
Most Canca1e exons are well-conserved throughout the vertebrate phylogeny. The UCSC genome browser provides base-specific Phastcons scores. For each exon of Cacna1e, we annotated the average Phastcons score.

## Extended data

Supplementary File 1: FLAIR generated FASTA file of all Cacna1e splice variants

Supplementary File 2: FLAIR generated counts table for all Canca1e splice variants

Supplementary File 3: Transdecoder predicted amino acid sequences for all splice variants

Supplementary File 4: All Cacna1e splice variants with a predicted CDD domain

Supplementary File 5: 154 Cacna1e splice variants with 4 pore-forming domains (predicted by CDD)

Supplementary File 6: Full BLAST results of Cacna1e splice variants

Supplementary File 7: Visualization of all 154 splice variants predicted to have 4 pore-forming domains

## Data availability

Raw long-read and short-read data will be submitted to GEO and SRA after manuscript acceptance.

## Author contributions

SB, JRT, TS and PP designed research. SB and JRT performed research. JS and MB contributed analytic tools. SB, JRT, TS and PP wrote paper.

## Acknowledgements

The authors would like to acknowledge the FLAIR developers for their help with their software. The authors would also like to thank members of the Pavlidis lab and the Snutch lab for their advice throughout the project.

