## Supplementary figures and images for "Cataloging the potential functional diversity of Cacna1e splice variants using long-read sequencing"

### Supplementary_file_7.pdf

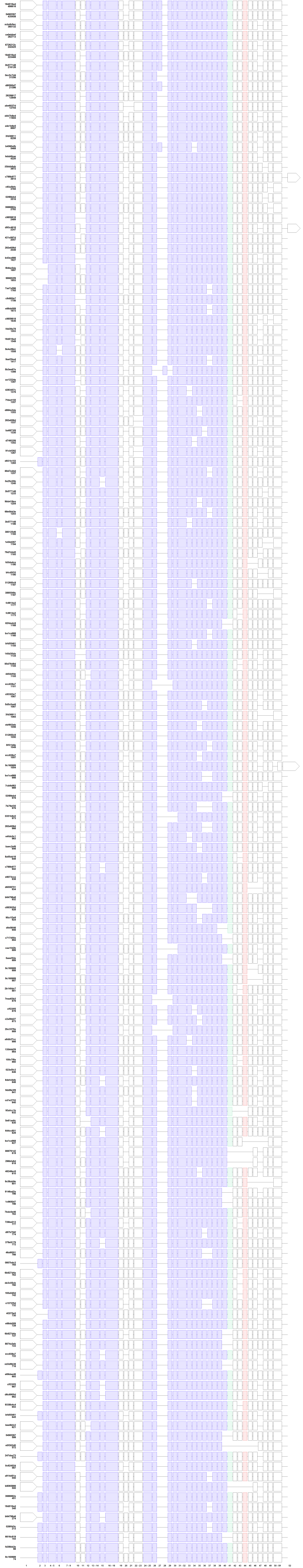
